# New semi-automated tool for the quantitation of MR imaging to estimate in vivo muscle disease severity in mice

**DOI:** 10.1101/2023.05.23.541310

**Authors:** Emily A. Waters, Chad R. Haney, Lauren. A Vaught, Elizabeth M. McNally, Alexis R. Demonbreun

**Author notes:** To whom correspondence should be addressed: Alexis R. Demonbreun PhD, Center for Genetic Medicine Northwestern University, 303 E Superior SQ 5-512, Chicago, Il 60611 USA.

## Abstract

The pathology in Duchenne muscular dystrophy (DMD) is characterized by degenerating muscle fibers, inflammation, fibro-fatty infiltrate, and edema, and these pathological processes replace normal healthy muscle tissue. The *mdx* mouse model is one of the most commonly used preclinical models to study DMD. Mounting evidence has emerged illustrating that muscle disease progression varies considerably in *mdx* mice, with inter-animal differences as well as intra-muscular differences in pathology in individual *mdx* mice. This variation is important to consider when conducting assessments of drug efficacy and in longitudinal studies. Magnetic resonance imaging (MRI) is a non-invasive method that can be used qualitatively or quantitatively to measure muscle disease progression in the clinic and in preclinical models. Although MR imaging is highly sensitive, image acquisition and analysis can be time intensive. The purpose of this study was to develop a semi-automated muscle segmentation and quantitation pipeline that can quickly and accurately estimate muscle disease severity in mice. Herein, we show that the newly developed segmentation tool accurately divides muscle. We show that measures of skew and interdecile range based on segmentation sufficiently estimate muscle disease severity in healthy wildtype and diseased *mdx* mice. Moreover, the semi-automated pipeline reduced analysis time by nearly 10-fold. Use of this rapid, non-invasive, semi-automated MR imaging and analysis pipeline has the potential to transform preclinical studies, allowing for pre-screening of dystrophic mice prior to study enrollment to ensure more uniform muscle disease pathology across treatment groups, improving study outcomes.

## Introduction

Duchenne muscular dystrophy (DMD) is an inherited muscle wasting disease caused by loss of dystrophin. Dystrophin, a large protein that links the cytoskeleton with the extracellular matrix, is encoded by the *DMD* gene. In the absence of dystrophin, the muscle membrane becomes easily disrupted which causes muscles to degenerate, and in its place, the intact muscle becomes replaced by fibrosis and fat, as well as immune infiltrate. The dystrophin-deficient *mdx* mouse model is routinely used in preclinical therapeutic trials as it recapitulates many of the same pathological features seen in humans with DMD. The *mdx* mouse model on the C57Bl/10 background harbors a naturally occurring mutation in exon 23 of dystrophin resulting in a premature stop codon ^1, 2^. Although the *mdx* mouse does not produce any detectable dystrophin protein, this model exhibits only mild histological and functional deficits in comparison to humans with DMD. A more severe mouse model referred to as *mdxD2* was generated through backcrossing the *mdx*C57Bl/10 model over 5 generations onto the DBA/2J strain ^3^. The muscle pathology in *mdxD2* mice has more similar features to human DMD pathology with increased areas of muscle damage, fibrofatty deposition, impaired repair capacity, and, along with these findings, greater functional deficits compared to the *mdx*C57Bl/10 model ^4-8^. The increase in disease progression on the DBA/2J background has been linked to multiple genetic modifiers, including Latent TGFB binding protein 4 (*Ltpb4*), annexin A6 (*Anxa6*), and osteopontin (*Spp1*) ^9-11^.

The availability of multiple mouse models of DMD has allowed for extensive preclinical studies evaluating potential therapeutics for the treatment of DMD. With preclinical testing occurring across multiple animal colonies, there have been efforts to standardize procedures to increase the reliability and translatability of endpoint measures (TREAT-NMD.org protocols). It is well known that the *mdx* model displays a wide range of both intra-animal and inter-animal variability ^12^. For example, *mdx* littermates can exhibit virtually no muscle damage or fibrosis in one subject, while another is severely affected. Moreover, within the same animal one muscle may have extensive pathology while the contralateral muscle is minimally affected. This inherent variation can contribute to poor assay sensitivity and necessitate large cohorts of animals to achieve the statistical power required to show treatment effects ^13^. Spurney et al showed that quantitative histology measures of degeneration and regeneration and creatine kinase showed high variance in the *mdx* model. Evans’ blue dye uptake, as a measure of muscle membrane leak, also showed both animal to animal variability and intra-animal variability across studies ^14^.

Understanding and mitigating factors that can influence preclinical outcome measures is critical when designing and evaluating therapeutic efficacy studies. Environmental factors such as cage design, light/dark cycle, food, sex, age, and weight can be more easily controlled than biological factors. Non-invasive imaging techniques such as magnetic resonance imaging (MRI) can provide information on *in vivo* muscle tissue health without the need for tissue sampling or sacrifice. Although MRI can accurately distinguish between healthy and diseased tissue, qualitative estimates of severity are highly subjective, and quantitative image analysis can be cumbersome, making it less than ideal as a screening tool. Further, it can be challenging to identify a numerical summary statistic that reflects qualitative differences readily observed between imaging datasets. For example, in datasets with small, localized regions of signal change, simple signal averaging can obscure differences due to the much larger areas of unchanged tissue. Indeed, previous efforts to use average T_2_ values as an imaging biomarker of disease progression in mdx mice have been hampered by high variance that has been interpreted as a lack of significant difference between timepoints or groups, but more likely reflects the inadequacy of bulk averaging as a summary statistic^15^. We propose a rapid high-throughput screening method that uses a combination of MR imaging, semi-automated segmentation, and quantitative analysis with meaningful summary statistics to estimate disease severity *in vivo*, balance treatment groups, and *a priori* exclude animals with unusually high or low disease burdens.

## Methods

### Animals

Male mice, 6-7 weeks old, WT, *mdxB10* and *mdxD2* mice were purchased from the Jackson Laboratory (stock 000664, 001801, and 013141). Mice were housed in a specific pathogen–free facility on a 12-hour light/12-hour dark cycle and fed ad libitum in accordance with the Northwestern University’s Institutional Animal Care and Use Committee regulations.

### MR Image acquisition

Mice were anesthetized in an induction chamber with inhaled isoflurane and then transferred to a dedicated imaging bed with a nosecone to deliver continuous isoflurane. Each mouse was positioned prone with legs tucked beneath the abdomen to reduce susceptibility artifacts. Respiration was monitored using a pressure sensitive pillow and warming was performed MRI was performed on a 9.4T Bruker Biospec 9430 (Bruker Corporation, Billerica, MA, USA) with a 30 cm bore and 12 cm gradient insert, running Paravision 6.0.1. Each mouse was in a 40 mm quadrature volume coil (Bruker) operating in transmit/receive mode. After localizer images were acquired, a T_2_ map was acquired using a spin echo sequence (Multi Slice Multi Echo, MSME) oriented axially and centered at the mid-calf. The following parameters were used: TR = 4000 ms, TE = 9-225 ms (30 echoes, echo spacing = 9 ms), MTX = 256 × 256, FOV 3.5 × 3.5 cm, 5 slices, 1 mm slice thickness and 1 signal average. Acquisition time was approximately 18 minutes.^16^

### Training of Segmentation Network

Images were imported into Amira 2020.2 software (Thermo Fisher Scientific, Waltham, MA, USA) and the first echo of the T_2_ map acquisition (TE = 9 ms), a relatively proton-density weighted image, was used to segment a region of interest (ROI) containing hindlimb and paraspinal muscles, and remove other tissues such as bladder, skin, fat, and chemical shift artifacts. As this was a time-consuming manual process, a deep learning prediction model was trained in Amira using the built-in tools. A training dataset was assembled using segmentation and image data 24 MDX scans and 7 wildtype scans and concatenated with a validation dataset assembled from segmentation and image data obtained by rescanning 5 MDX mice and 3 wildtype mice 2-3 days after their initial imaging session. A preliminary model was trained using the “Deep Learning Training” module in Amira with type BackbonedUNet, number of classes = 2, resnet18 backbone, patch size 128, batch size 8, max patch overlap 0.3, maximum number of epochs 100, Adam optimizer, learning rate 0.0001, and no data augmentation. A refined model was trained using the same module, initialized with the weights of the preliminary model, with the following modifications: patch size 256, maximum number of epochs 500, learning rate 0.0005, and data augmentation with geometry transforms including horizontal and vertical flip, 30-degree rotation, 10% zoom, and 10 degree shear, and an early stopping criterion to stop if no model improvement for 25 iterations. The “Segmentation Metrics” Amira Xtra module was used to assess the performance of the model on the validation dataset.^17^

### Semiautomated Image Segmentation

After the deep learning model was generated, segmentation was performed by importing the first echo of the T_2_ map acquisition into Amira and generating a segmentation using the “Deep Learning Prediction” module using the established model weights. The segmentation was inspected by a trained observer and manually refined as necessary to correct minor errors (e.g., incomplete removal of the bladder). Muscle ROIs were exported from Amira as a mask image to be used in further processing steps.

### MR Image Processing

All echoes of the T_2_ map acquisition, were imported into JIM 7 (Xinapse Systems Ltd, West Bergholt, Essex, UK), masked to include only muscle voxels using the mask image generated in the segmentation step, and fit using the built in nonlinear curve fitting module. A custom Python script was used to extract a list of muscle T_2_ values from the resultant maps. T_2_ values < 15 ms and > 100 ms were excluded as these overwhelmingly corresponded to voxels with low signal and low-quality fitting.

### Evans Blue Dye

To validate the skew and interdecile range of the T_2_ distribution as metrics for disease severity, they were compared to Evans Blue dye uptake. WT mice (n=5), *mdx* mice (n=4) and *mdxD2* mice (n=3) underwent the MRI protocol described above and then underwent measurements of Evans Blue dye uptake that were quantified as described previously ^9, 18, 19^. Briefly, mice were injected with 5 µl per gram of body weight of 10 µM Evans Blue dye (E2129, Sigma-Aldrich). Mice were euthanized, tissue excised, minced, and placed in 1ml of formamide in a 24-well plate placed at 55 °C for 2 hrs. Subsequently, absorbance was measured at 620 nm on a Synergy HTX multi-mode plate reader (BioTek®) from 200 µl of sample solution in a 96-well plate. Each sample was assessed in duplicate. Results are reported as the average arbitrary optical density units/mg of tissue.

### Statistical Analysis

A key purpose of this study was to identify and validate relevant summary statistics for MRI image distributions based on the hypothesis that the distribution of T_2_ values would be sufficiently non-normal as to render mean and standard deviation inappropriate summary statistics. A custom Python script was used to test the normality of the distribution of T_2_ values for each animal using D’Agostino’s K2 test, and compute summary statistics.^20-24^ All distributions observed were markedly non-normal (p < 0.001). As a result, summary statistics of skew and interdecile range were calculated to describe the width and asymmetry of distributions and enable quantitative comparisons. Correlation coefficients for the test-retest variability and the comparisons of skew and IDR with Evans’ Blue dye uptake were calculated using a simple linear regression model.

### Code Availability

Imaging data and associated code for processing and analysis is available online.^25^

## Results

### Development of a semi-automated muscle segmentation tool

Manual segmentation of the muscle ROI including the hindlimb and paraspinal muscles was a lengthy manual process due to the need to manually remove bone signals as well as signals from normal fluid and fat that are present subcutaneously and at intramuscular interfaces. The signals from regions of fat additionally required removal of chemical shift artifacts. Because of the manual and subjective nature of this process, it was highly subject to intra- and interoperator variability. Use of the deep learning model reduced the time for generating muscle ROIs from over an hour to a few seconds and reduced total hands-on analyst time to 5-10 minutes. In the validation dataset, the model-predicted label field matched the ground truth label field with a Jaccard Index of 0.97, indicating excellent agreement. An example segmentation of a dataset not included in the training data is shown in **Figure 1**.

**Figure 1:**
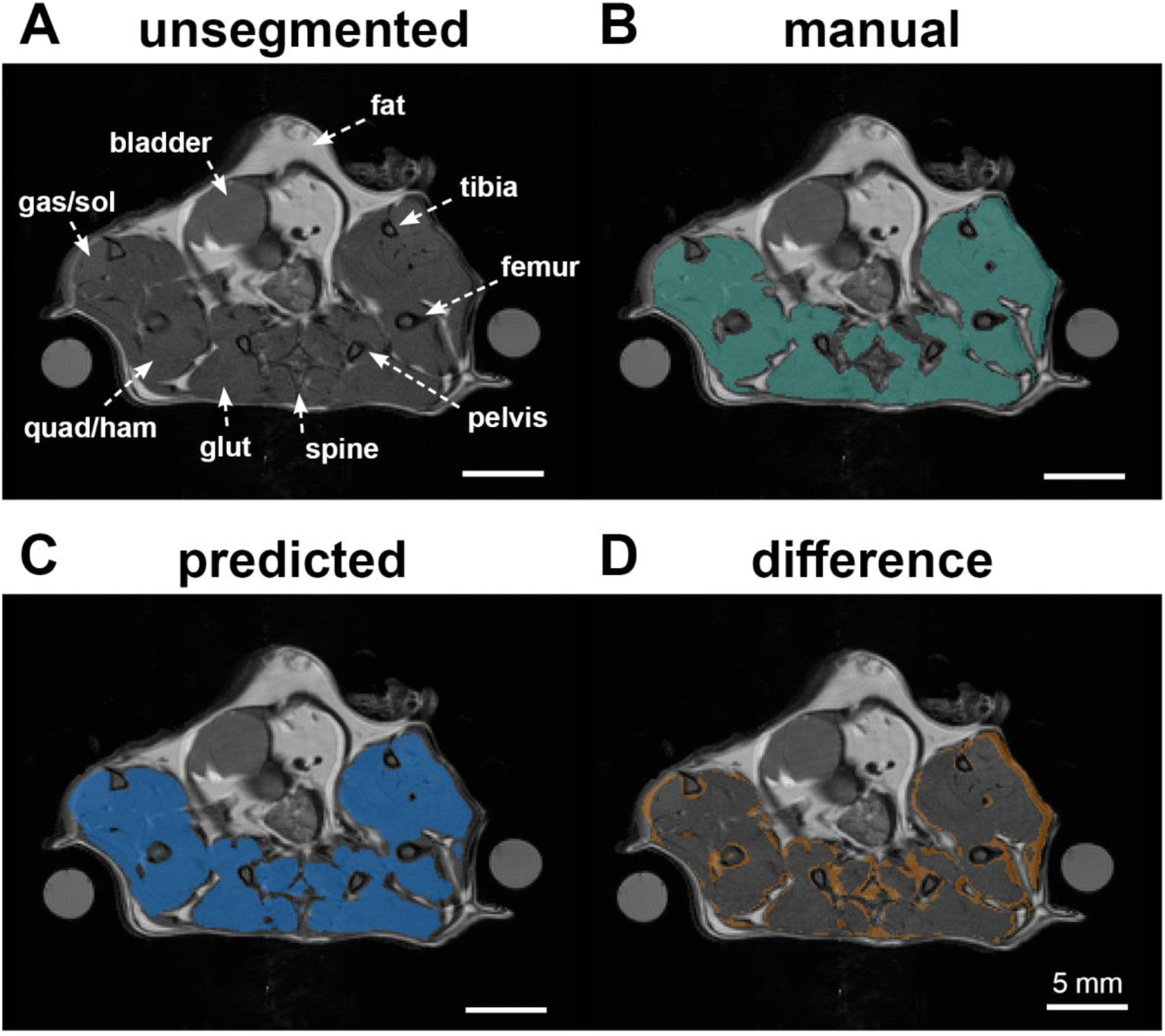
Reliable semi-automatic muscle segmentation. (**A**) Proton density weighted axial image showing hindlimb and paraspinal musculature. Muscle groups visualized include gastrocnemius and soleus (lower hindlimb), quadriceps and hamstrings (upper hindlimb) and gluteal muscles (near pelvis). (**B**) Manual segmentation of muscle ROI. (**C**) Result of segmentation using the trained deep learning model. (D) Difference between manual and automated segmentation showing overall excellent agreement with minor variation around the muscle-skin and muscle-bone interfaces.

### Stratification of muscle disease through semi-automated MR imaging analysis

*mdxB10* mice expressing a severe disease phenotype (**Figure 2A**), mild disease phenotype (**Figure 2B**), and healthy wildtype controls (**Figure 2C**) were imaged to cover a wide range of muscle health. Examination of the T_2_ maps shown in **Figure 2A** and **Figure 2B** illustrates the wide range of disease phenotype present in this cohort of same-aged and same-sex mdx mice. **Figure 2A** illustrates a mouse with severe disease as indicated by extensive regions of mildly elevated T_2_, in addition to several focal regions of highly elevated T_2_. In contrast, **Figure 2B** illustrates a mouse with comparatively mild disease, in which there are focal regions with mildly elevated T_2_, and many fewer voxels with highly elevated T_2_. The healthy wildtype mouse shown in **Figure 2C** has few voxels with high T_2_ and these are primarily located at the intermuscular interfaces. For each T_2_ map, the distribution of T_2_ values is plotted immediately to the right, with the mean plotted in red and the standard deviation represented by red error bars (**Figure 2A-C**). The distribution of T_2_ values is immediately seen to be non-normal (as confirmed by D’Agostino’s K^2^ test of normality). A plot of skew vs. interdecile range of the distribution of T_2_ values was effective in stratifying disease severity and correlated well with visual assessment of the selected T_2_ maps **(Figure 2D)**. Together these data show that MR image-based measures extracted using the newly developed, semi-automated segmentation and analysis pipeline stratify muscle disease severity across healthy to severe pathology.

**Figure 2:**
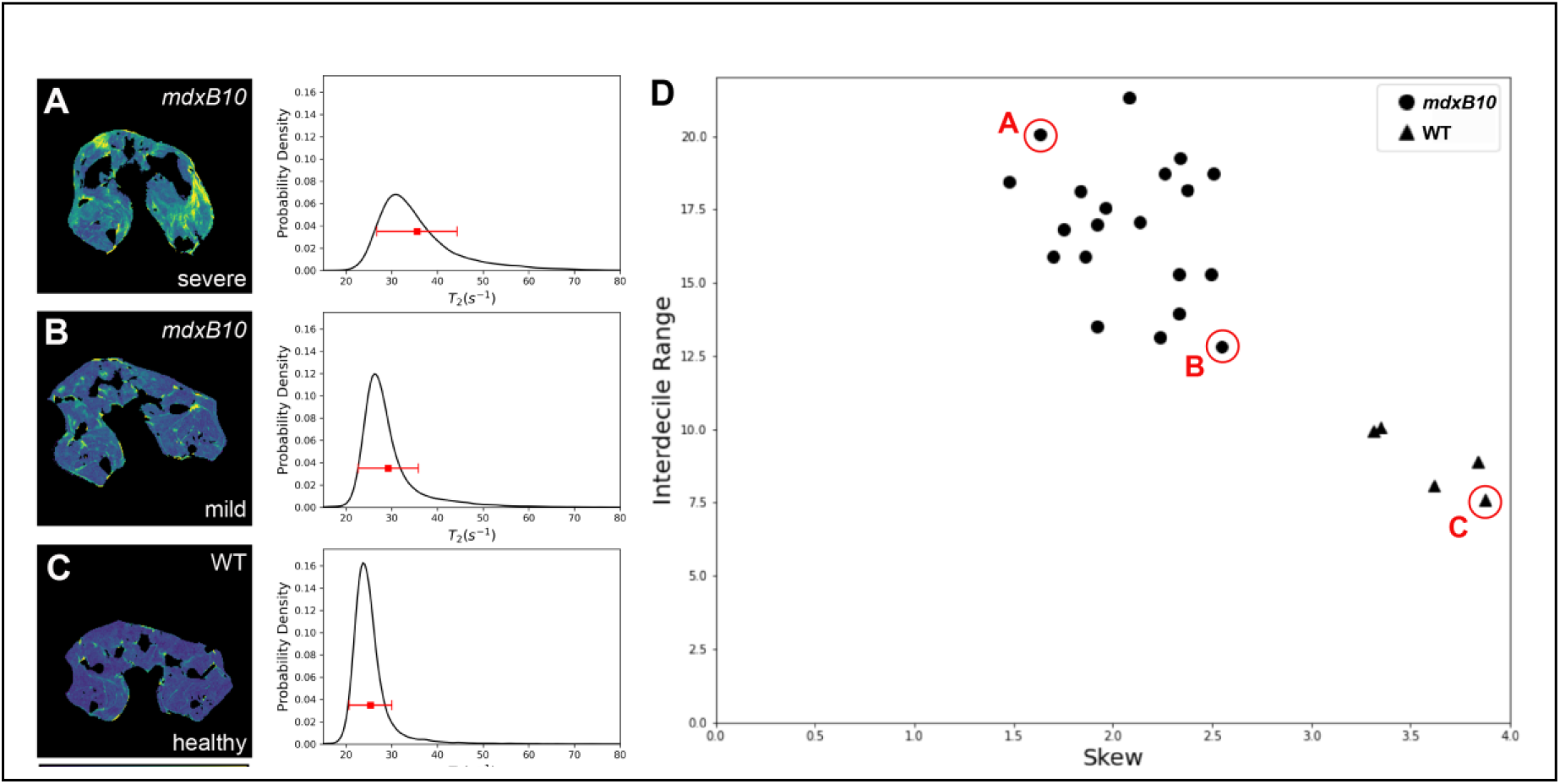
MR image-based measures stratify muscle disease severity. (**A-C**) on the left are representative T_2_ map slices from 3 different animals in the study. (**A**) shows an *mdxB10* animal with a severe muscle disease phenotype as evidenced by the globally higher T_2_ values as well as localized areas of high T_2_ corresponding to muscle damage. (**B**) shows an *mdxB10* animal with a relatively mild disease phenotype seen by moderate regions of high T_2_ values. (**C**) is a healthy wildtype (WT) control animal showing minimal regions of high T_2_ aside from normal intramuscular interfaces. Plots to the right of each image show the distribution of T_2_ values for that animal. The red square and error bars correspond to the mean and standard deviation of the T_2_ values and demonstrate the insensitivity of those metrics to changes in the shape of the distribution. (**D**) A plot of skew vs interdecile range (IDR) for 20 *mdxB10* animals and 5 healthy WT control animals, demonstrating that skew and IDR clearly separate the two genotypes as well as stratifying disease severity for the *mdx* animals. Data points marked A-C correspond to the individual datasets shown on the left.

### Semi-automated MR image-based measures are reproducible across disease severity

To assess the repeatability of measurements of the skew and interdecile range of the T_2_ distribution, 5 *mdx* mice and 3 WT mice were reimaged two days after their initial imaging session. Both sets of data were analyzed by a trained observer according to the protocol described above. Both skew and interdecile range were found to be highly repeatable across imaging sessions (**Figure 3**). The first and second measurements were highly correlated for both skew (R^2^ = 0.97, p < 0.001) and interdecile range (IDR) (R^2^ = 0.81, p = 0.002)

**Figure 3:**
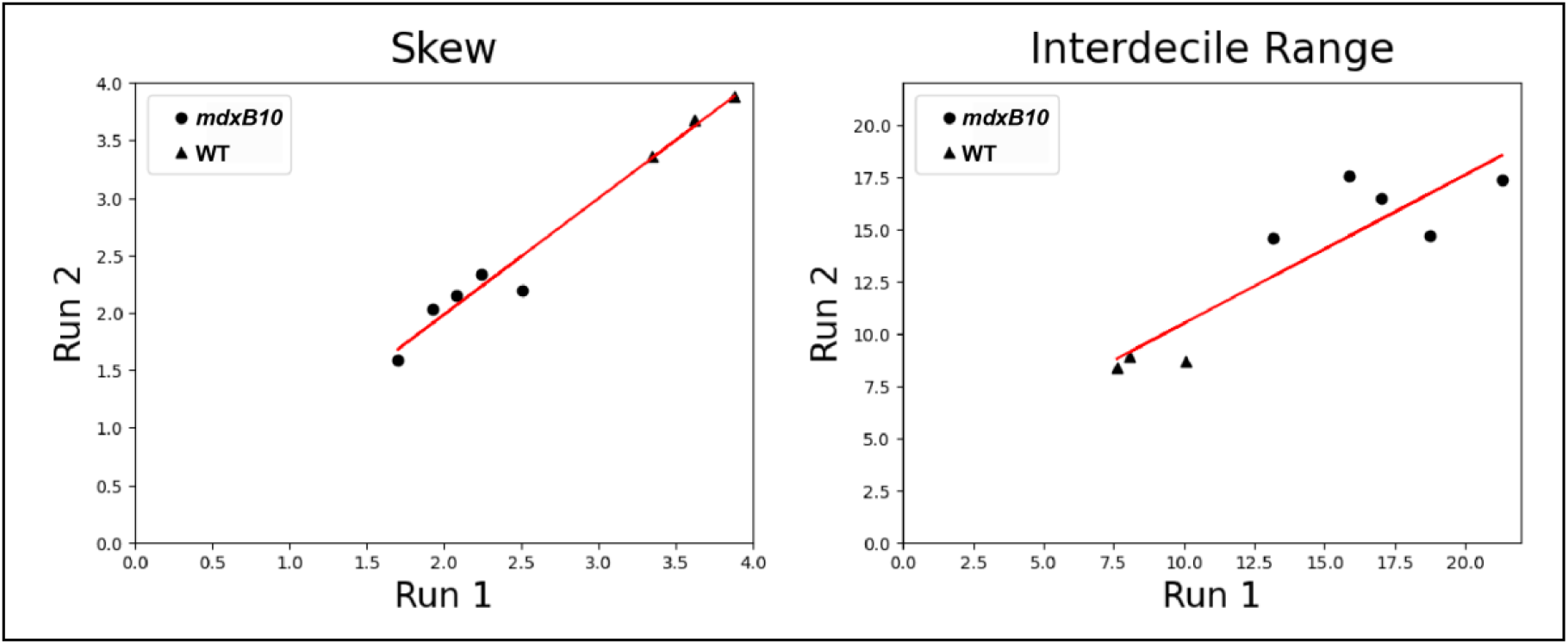
Reproducible MR image-based measures in healthy and diseased muscle. Test-retest reliability comparison of measured values of skew (left) and IDR (right) in *mdxB10* mice reimaged 2 days after the initial imaging session. Both values exhibited a strong linear correlation with skew R^2^ = 0.97 and IDR R^2^ = 0.81 including diseased (mdx) and healthy (WT) muscle.

### High correlation between measures of muscle membrane leak and semi-automated MR imaging-based measures

Quantification of Evans blue dye uptake is commonly used to evaluate muscle membrane permeability and stability as a measure of muscle health, as dye is excluded from healthy intact muscle. To determine if the MR image-based measures, skew, and IDR, correlate with dye uptake measures, *mdxB10, mdxD2*, and WT mice underwent MR imaging and subsequently were evaluated for Evans blue dye uptake in hindlimb muscles. Good agreement was observed between the *in vivo* imaging based metrics (skew and IDR) and the *ex vivo* Evans Blue dye uptake measurements (**Figure 4**). Healthy WT muscle had the lowest level of dye uptake and IDR followed by *mdxB10*, with *mdxD2* muscle taking up the highest levels of dye and having the highest IDR values. IDR exhibited a strong linear correlation with the Evans Blue Dye with R^2^ = 0.9 (p < 0.001). Skew was also linearly correlated with R^2^ = 0.71 (p < 0.001). This suggests that the *in vivo* imaging based metrics can serve as a surrogate measure of muscle membrane health and integrity as compared to the terminal Evans blue dye uptake assessment.

**Figure 4:**
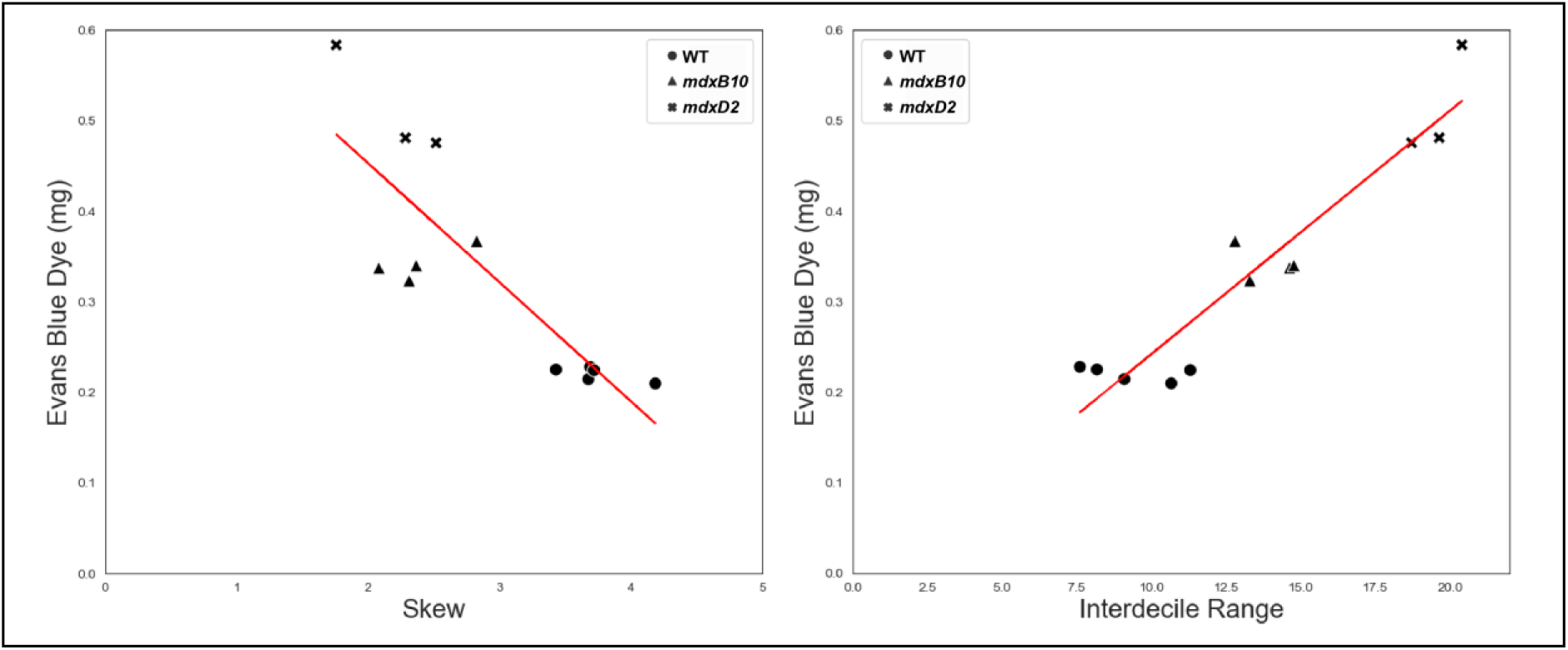
Strong correlation between dye uptake and MRI measures across muscle disease severity. Comparison of noninvasive *in vivo* image-based measurements of disease severity (skew and interdecile range) with *ex vivo* measurement of Evans Blue Dye. No muscle disease occurred in healthy WT controls (n=5), mild disease was present in *mdxB10* mice (n=4), and severe disease was evident in *mdxD2* mice (n=3). Both measures exhibited a strong linear correlation with the Evans Blue Dye (R^2^ vs skew: 0.71; R^2^ vs IDR: 0.9).

## Discussion

We have outlined a non-invasive, quick, and reproducible imaging and analysis protocol that can accurately stratify animals to aid in preclinical therapeutic study design. Screening of animals to assess disease severity prior to assigning treatment groups can help mitigate the potential effects of the wide range of disease severities in the *mdx* model, as treatment effects can be obscured if there are differences in the underlying severity of disease between the groups. Many studies rely on comparing histopathological findings or Evans Blue dye uptake in separate groups of untreated and treated animals. These measures rely on terminal procedures that cannot be undertaken at the outset of a study.

Previous efforts to use MRI in the assessment of disease severity in *mdx* mice have focused on T_2_ maps and T_2_ weighted images because regions of muscle damage tend to have abnormally elevated T_2_ values. However, researchers have consistently noted very small differences in the average value of muscle T_2_ (on the order of a few milliseconds) between *mdx* and wildtype animals.^15, 26^ Such a small difference, while in some cases statistically significant, can be easily confounded by minor systemic errors in image acquisition and fitting. Upon deeper interrogation of our study data, we found that the mean and standard deviation of T_2_ values over a region of interest comprising hindlimb and paraspinal musculature were very poor descriptors of the actual distribution of measured T_2_ values and were not appropriate for stratifying the severity of disease in *mdx* animals. Attempts to differentiate the groups by setting a T_2_ threshold and counting voxels above that threshold were highly dependent on the specific threshold selected and ineffective at stratifying disease.

Inspection of the distribution of T_2_ values for a muscle region of interest in *mdx* and WT mice yielded stark differences in the peak shape and width of the distribution of T_2_ values between the *mdx* and WT mice. In particular, in WT mice, the distribution of T_2_ values was a sharp narrow peak roughly centered around a low T_2_ value, with a shallow tail of higher values corresponding to blood vessels, inter-muscular interfaces, and noisy voxels with poor T_2_ fitting. In contrast, for the *mdx* mice, the distribution was shifted rightwards in the direction of higher T_2_ values, and markedly asymmetric with a “heavy” tail of high T_2_ values, mostly corresponding to regions of edema and necrosis. The distributions were assessed for normality and found to be markedly non-normal, confirming that the mean and standard deviation were inappropriate summary statistics for these data.

Because the goal was to identify animals with a higher concentration of high-T_2_ voxels, corresponding to the heaviness of the distribution tail, skew was investigated as a higher-order summary statistic and interdecile range as a measure of spread of the values, on the premise that these metrics would better reflect the shape of the distribution and might be more suitable for stratifying degree of disease. The plot of these two metrics correlated with the spectrum of disease severity, clearly and repeatably differentiating *mdx* animals from wildtype animals and correlating well with the terminal measurements made using Evans Blue Dye. In this study, skew and IDR were identified as metrics suitable to describe the acquired data. In future studies, this analysis will be extended to a more comprehensive radiomics approach in an effort to identify texture-based image features that may reflect localized areas of muscle damage.^27^

A major strength of this rapid semiautomated imaging and analysis pipeline is that images of several major hindlimb muscle groups can be acquired in approximately 30 minutes, with only 10-15 minutes per animal needed for semiautomated segmentation and processing. This significantly increases the throughput relative to manual segmentation and processing, which required 1.5-2 hours per animal, and renders it suitable for screening of cohorts of animals prior to assigning treatment groups. The proposed pipeline could substantially improve the quality of studies of therapeutic efficacy in mouse models of muscular dystrophy by ensuring that study cohorts are appropriately balanced across the spectrum of disease and a priori excluding animals with either extremely severe or extremely mild disease.

## Abbreviations

(DMD): Duchenne Muscular Dystrophy
(MRI): magnetic resonance imaging

## AUTHOR CONTRIBUTIONS

LV and AD performed the mouse harvest and EBD study. EW and CH performed the MR imaging, segmentation, and analysis. EW, CH, AD and EM conceived and designed the studies, analyzed the data, and wrote the manuscript.

## ACKNOWLEDGEMENTS

This work was supported by National Institutes of Health AR052646, HL167813, NS047726, NS127383, Additional funding was through Lakeside Discovery. MRI was performed at the Northwestern University Center for Advanced Molecular Imaging (RRID:SCR_021192) generously supported by NCI CCSG P30 CA060553 awarded to the Robert H Lurie Comprehensive Cancer Center. EAW is supported by grant number 2020-225578 from the Chan Zuckerberg Initiative DAF, an advised fund of Silicon Valley Community Foundation.

